# Similar visual perception in GCaMP6 transgenic mice despite differences in learning and motivation

**DOI:** 10.1101/2020.02.18.954990

**Authors:** Peter A. Groblewski, Douglas R. Ollerenshaw, Justin Kiggins, Marina Garrett, Chris Mochizuki, Linzy Casal, Sissy Cross, Kyla Mace, Jackie Swapp, Sahar Manavi, Derric Williams, Stefan Mihalas, Shawn R. Olsen

## Abstract

To study mechanisms of perception and cognition, neural measurements must be made during behavior. A goal of the *Allen Brain Observatory* is to map activity in distinct cortical cell classes during visual processing and behavior. Here we characterize learning and performance of five GCaMP6-expressing transgenic lines trained on a visual change detection task. We used automated training procedures to facilitate comparisons across mice. Training times varied, but most transgenic mice learned the task. Motivation levels also varied across mice. To compare mice in similar motivational states we subdivided sessions into over-, under-, and optimally motivated periods. When motivated, the pattern of perceptual decisions were highly correlated across transgenic lines, although overall d-prime was lower in one line labeling somatostatin inhibitory cells. These results provide important context for using these mice to map neural activity underlying perception and behavior.

## Introduction

Goal-oriented behavior requires coordinated neural activity across brain regions, but the cellular mechanisms mediating these activity dynamics are not fully understood. The mouse provides unique opportunities to dissect cell type- and circuit-specific mechanisms of perception and behavior (Luo et al., 2018, 2008; Niell, 2015). Head-fixed behaviors are well-established and allow precise measurements of cellular activity using 2-photon imaging and electrode recordings, in addition to optogenetic perturbations (Andermann et al., 2010; Burgess et al., 2017; Z. V. Guo et al., 2014; Histed et al., 2012; O’Connor et al., 2010). Applications of these methods are revealing mechanisms of perception and action across multiple sensory modalities and cognitive systems (Chen et al., 2013; Glickfeld et al., 2013; Goard et al., 2016; Z. Guo et al., 2014; Harvey et al., 2012; Huber et al., 2012; Li et al., 2016; O’Connor et al., 2013; Peron et al., 2015; Petreanu et al., 2012; Pinto et al., 2013; Poort et al., 2015; Resulaj et al., 2018).

At the Allen Institute for Brain Science we seek to generate a database of cell type-specific activity across visual cortical areas during visual stimulation and behavior (Koch and Reid, 2012). We developed a standardized physiological pipeline—the *Allen Brain Observatory*—to monitor cellular population activity during passive visual stimulation in mice (de Vries et al., 2019). To expand on these passive viewing datasets, we are adapting our existing pipeline to include recordings from mice performing visually-guided behaviors. For large-scale pipeline compatibility we seek tasks that are simple yet adaptable to more complex variants, easily learned, and consistently performed. Candidate tasks must also support head-fixed physiological measurements using our standardized instruments.

In this study we test a go/no-go visual change detection task. Change detection is a fundamental behavioral capacity of animals and humans (Elmore et al., 2011; Hagmann and Cook, 2013; Pearson and Platt, 2013; Rensink, 2002), and the visual cortex of mice and primates is implicated in the detection of changes in visual features (Brunet et al., 2014; Glickfeld et al., 2013; Womelsdorf et al., 2006). The core task we use can be used to test perception of various visual features including orientation, contrast, color, and natural images (Denman et al., 2018; Garrett et al., 2020; Glickfeld et al., 2013). Moreover, our task includes features that permit investigation of the physiological correlates of behavior and cognition. For instance, the ability of mice to generalize to new stimuli allows for exploration of stimulus novelty and learning, and the regular temporal structure of the task allows for exploration of deviations from expected timing (Garrett et al., 2020). Additionally, the delay between stimulus presentations provides a test of short-term memory. Finally, variability in task-engagement and motivation provides a window into state-dependent processing.

To support future studies of neurophysiology during this versatile task, we characterize the behavior of five Cre driver x GCaMP6 reporter transgenic mouse lines that label subpopulations of excitatory or inhibitory cells—these allow cell class-specific activity mapping (de Vries et al., 2020; Garrett et al., 2020; Madisen et al., 2015). To mitigate sources of variability in behavior and facilitate inter-mouse comparisons we used automated training procedures to progress mice through a series of increasingly difficult training stages.

Even in well-trained subjects, psychophysical performance can be non-stationary over a behavioral session, varying with motivation, attention, confusion, and other factors (Andermann et al., 2010; Berditchevskaia et al., 2016; Carandini and Churchland, 2013; Mcginley et al., 2015). Tasks using water restriction, as in our study, are subject to motivational changes due to decreasing thirst as water is consumed during the session. Studies often only consider average performance over the session or restrict session duration to avoid major motivational changes. Here, all mice completed one-hour sessions, independent of mouse performance and experimenter intervention. Inspired by a recent study of motivation dynamics in mice performing a go/no-go task (Berditchevskaia et al., 2016), we use the signal detection theory metric, ‘criterion’, to help categorize epochs in the session as over-motivated, motivated, and under-motivated. Parsing behavior sessions according to motivation level helps to compare behavior and physiology across mice and transgenic lines under more controlled conditions.

Overall, each of the transgenic mouse lines we tested could be trained with automated algorithms to reach high performance levels, although training times varied across mice, and in some cases across lines. Additionally, we observed motivational and overall performance (d-prime) differences in some lines. However, we show that the pattern of perceptual decisions is highly correlated across transgenic mice during epochs of matched motivation. These results provide a basis for systematic neural activity mapping using these transgenic mice.

## Results

### Visual change detection task with natural scene images

We trained mice (n = 60) to perform a visual change detection task with natural scene images chosen from the *Allen Brain Observatory* battery of visual stimuli (http://observatory.brain-map.org/visualcoding). In this go/no-go task, mice see a continuous series of briefly presented images and they earn water rewards by correctly reporting when the identity changes (Figure 1). Responses are indicated by licking a water spout within a 600 ms response window following the image change (Figure 1A,B). On randomly interleaved ‘catch’ trials, no image change occurs and the mouse must withhold licking to avoid a time-out (Figure 1A,B). Licks that came before the randomly selected change time on a given trial resulted in that trial being aborted, leading to a short timeout followed by a reset of the trial clock (see Supplementary Figures 1 and 2 for detailed task flow). Once trained, mice display short latency reaction times with the majority of responses occurring within the response window (Figure 1C).

**Figure 1.**
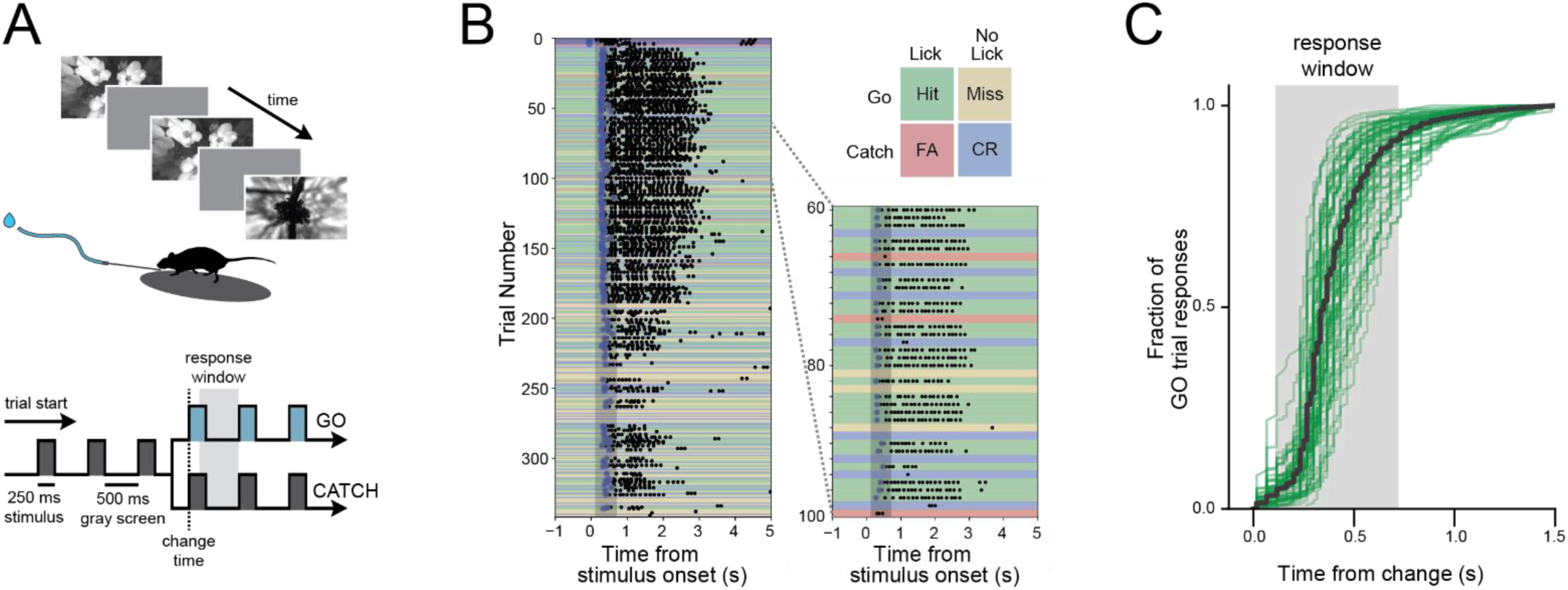
Change detection task with natural images. A) Behavioral task. Visual stimuli are shown for 250 ms with an intervening gray period of 500 ms. On GO trials, the image identity changes and mice must lick within the 600 ms response window to receive a water reward. On CATCH trials, no image change occurs, and licking is measured to quantify guessing behavior. B) Example of a complete behavior session with trials aligned to the time of image change. Trial types and outcomes are illustrated in the 2×2 matrix. C) Cumulative reaction time distribution on GO trials. Green lines show individual mice (n=56), and the black line indicates the average of all mice.

In our behavioral apparatus, mice are head-fixed yet free to run on a circular disc. Running is monitored but does not influence task flow. Most, but not all, mice ran or walked during the behavioral session, and these mice typically stopped running when responding to stimulus changes and to consume the water reward (Supplementary Figure 2).

### Automated behavior training of transgenic mice

We assessed training and performance of five transgenic mouse lines expressing GCaMP6f in distinct subsets of cortical cells (*Cux2*: Cux2-CreERT2;Camk2a-tTA;Ai93(TITL-GCaMP6f), n=4; *Rbp4*: Rbp4-Cre_KL100;Camk2a-tTA;Ai93(TITL-GCaMP6f), n=12; *Slc17a7*: Slc17a7-IRES2-Cre;Camk2a-tTA;Ai93(TITL-GCaMP6f), n=23; *Sst*: Sst-IRES-Cre;Ai148(TIT2L-GC6f-ICL-tTA2), n=7; *Vip*: Vip-IRES-Cre;Ai148(TIT2L-GC6f-ICL-tTA2), n=14). To train these transgenic mice (Cux2, Rbp4, Slc17a7, Sst, Vip) in a standardized manner, we developed an automated protocol in which mice progress through a series of training stages with parameters, performance requirements, and stage transitions defined in software rather than relying on experimenter intervention (Figure 2A; see Methods).

**Figure 2.**
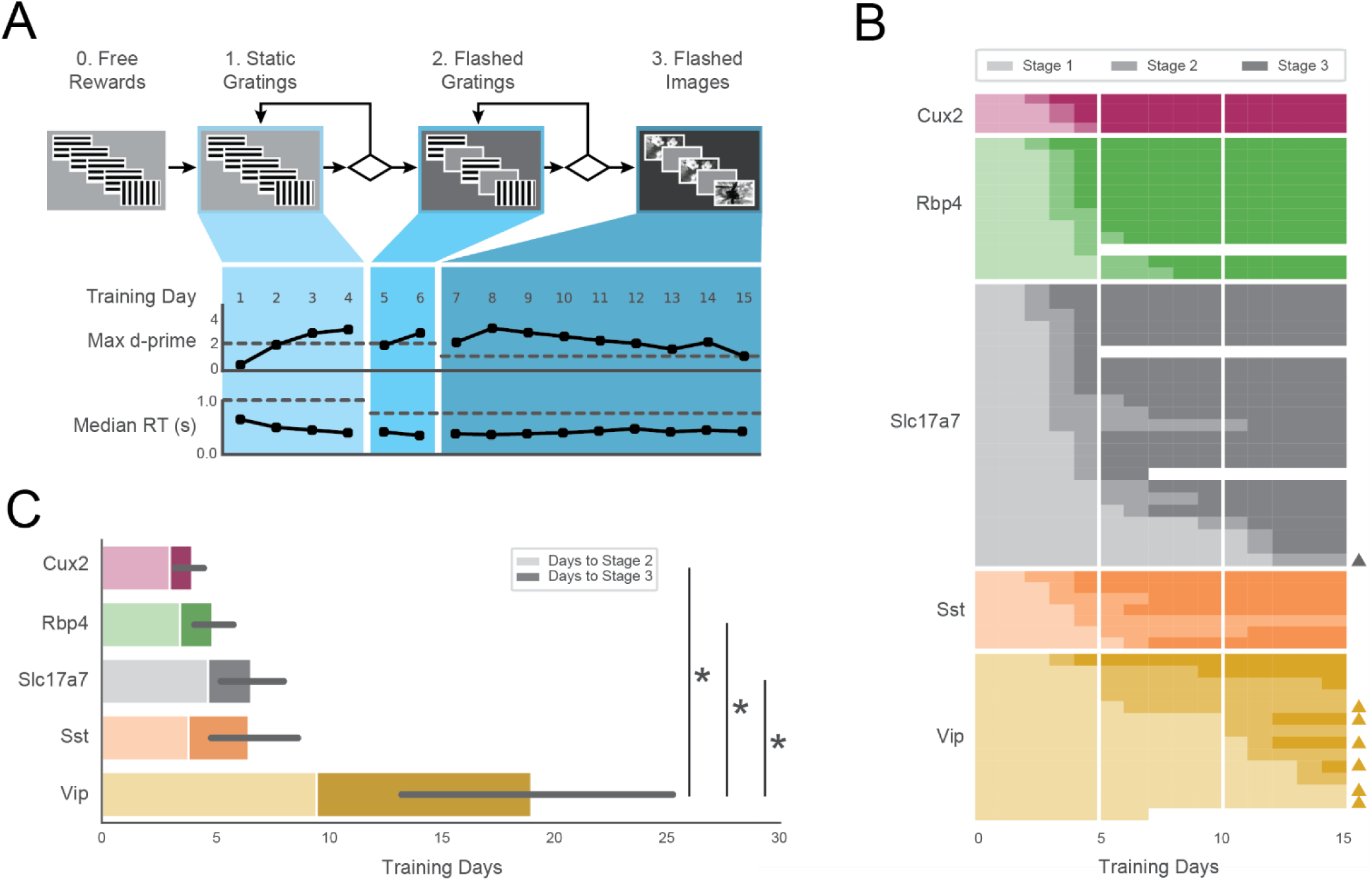
Automated training of five GCaMP6-expressing transgenic lines. A) *Top:* Progression of training stages. *Bottom:* Training trajectory for one example mouse (M328341, genotype: Rbp4). Max d-prime (in 100-trial rolling window) and median reaction time are shown for each training day. Horizontal dashed lines represent the max d-prime required for advancement and the maximum reaction time following stimulus change which would result in reward. B) Training days in each stage (each row is one mouse). Opacity of bar indicates training stages 1-3 in (A). Triangles on right indicate mice that reached Stage 3 after 3 weeks of training. Some mice (n=4) were removed from training early due to a health-related issue. C) Average number of sessions required to reach Stage 2 (light shading) and Stage 3 (dark shading) for all groups. Non-parametric analysis showed a significant main effect of group on time to Stage 3, with Vip mice exhibiting significantly longer training times than Slc17a7, Rbp4, and Cux2 mice.

Mice first learn the task with oriented gratings and no intervening gray period between stimuli. After reaching performance requirements on the orientation task, a 500 ms inter-stimulus gray period is introduced. In the final training stage, the grating stimuli are replaced with natural scene images. The majority of mice (47/60) completed the full set of training stages within 15 sessions, and 56/60 mice reached the final stage within 40 sessions (Figure 2B). The average time to reach the final training stage varied across genotypes (Figure 2C; Cux2, 4.0±0.8; Rbp4, 4.9±1.4; Slc 6.6±3.5; Sst, 6.5±2.6, Vip, 19.0±10.9), and there was a significant main effect of genotype on training times (H=22.98, *p*=0.0001). Post-hoc, pairwise comparisons showed Vip transgenic mice were slower to train than the Slc (*p*=0.0002), Rbp4 (*p*=0.0005), and Cux2 groups (*p*=0.003). Thus, all genotypes were able to learn the task, but the number of sessions to do so varied.

All subsequent data analysis is restricted to sessions in the final training stage (stage 3) in which mice had peak hit rate and d-prime values (both calculated over a rolling 100 trial window) of at least 0.3 and 1.0, respectively, and had at least 50 correct responses on hit trials. Of 1319 sessions in the final training stage, 1100 met these performance criteria. Of the 60 mice in the study, 56 mice had at least one stage 3 session (median = 21, mean = 19.7, standard deviation = 11.3, min=1, max = 37). Supplemental Table 1 provides a detailed summary of the mice described in this study, including the number of sessions analyzed.

### Variation in motivation

In typical behavior sessions, mice were very responsive early but became less task-engaged later in the hour-long session. During these periods of reduced task-engagement, mice licked only infrequently, or ceased licking altogether, indicating that motivation to perform the task decreased (Figure 3A).

**Figure 3.**
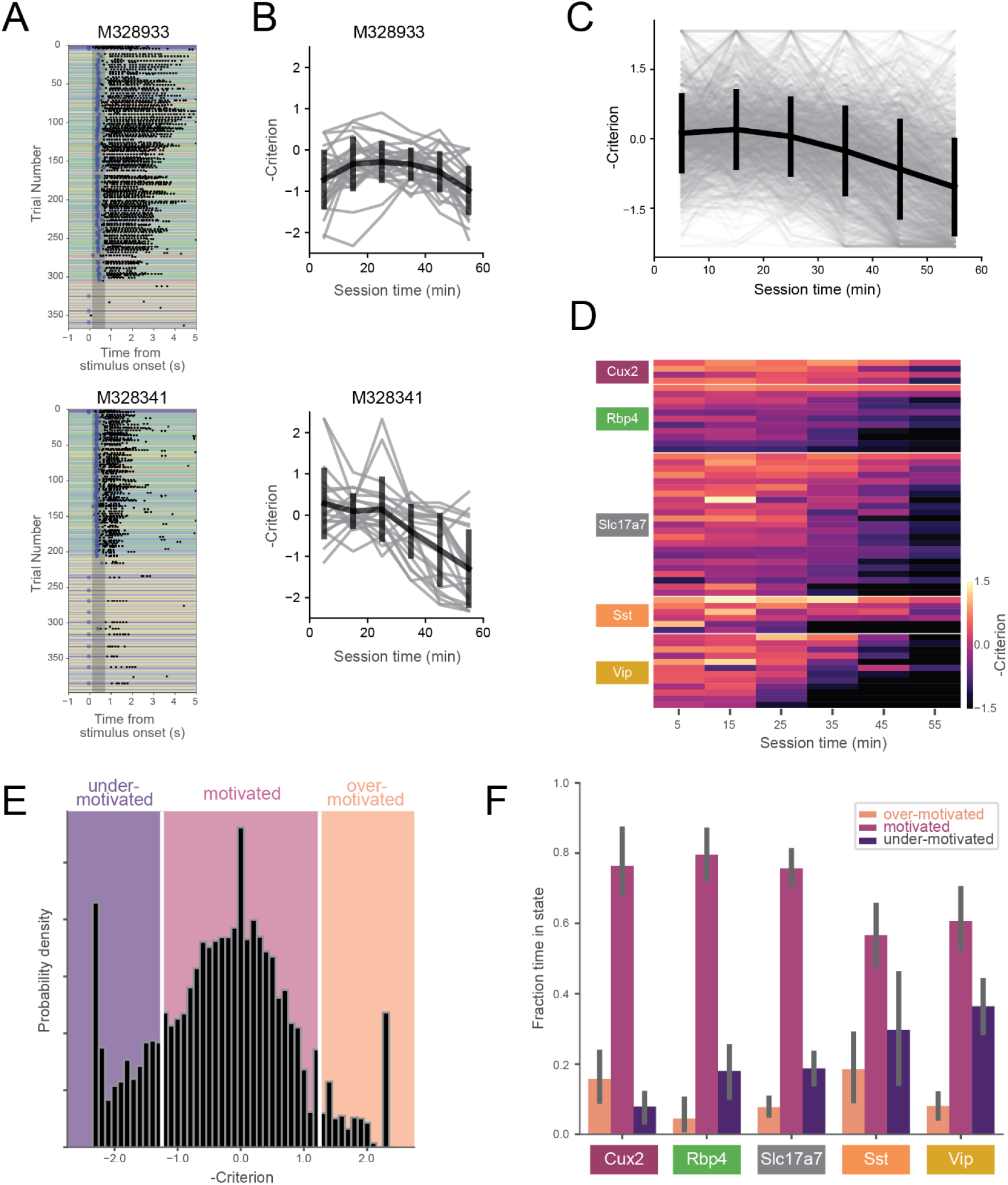
Motivation decreases over behavioral session. A) Example behavioral sessions from two mice showing high task-engagement early in the session followed by later disengagement. B) Criterion (−0.5*(z(HR)+z(FA))) computed in 10-minute bins for same mice in (A). Individual sessions are shown in gray and the mean over all sessions (+/-SD) is shown in black. C) Criterion in 10-minutes bins for all sessions (gray) and mean (+/-SD) across all mice (black). D) Across-session average of criterion values for all mice (each row represents a single mouse). E) Histogram of criterion values (10-minute epochs). White lines indicate boundaries for defining 3 motivation states: ‘over-motivated’ (criterion > 1.25), ‘motivated’ (−1.25 ≤ criterion ≤ 1.25), and ‘under-motivated’ (criterion < −1.25). F) Fraction of session epochs spent in each engagement state. All groups except Sst and Vip groups spent significantly more time in motivated versus under-motivated states.

We quantified changes in motivation using the ‘criterion’ parameter from signal detection theory (−0.5*[z(HR)+z(FA)]). Criterion is a measure of the subject’s internal bias to respond. Higher values correspond to more conservative response criteria and correspondingly lower response rates. To aid visualizations we represent criterion with the sign inverted, thus mapping states of low motivation to lower values and states of high motivation to higher values. To capture motivation changes over the course of the behavioral session, we computed criterion in ten-minute epochs. On average, mice showed decreasing motivation over the course of the one-hour session (Figure 3B,C), but we observed a range of motivation levels across mice and genotypes (Figure 3D).

To compare mouse behavior during similar motivational states, we subdivided behavioral sessions into epochs labeled ‘over motivated’ (criterion > 1.25), ‘motivated’ (−1.25 ≤ criterion ≤ 1.25), and ‘under motivated’ (criterion < −1.25) (Figure 3E). A small percentage of epochs (1.2%) were not assigned a criterion value due to insufficient presentations of GO and/or CATCH trials in 10-minute epoch (Supplemental Table 1). Mice spent the majority of their time in the ‘motivated’ state (Figure 3F), however, there was a significant interaction between genotype and state (F(8,102)=4.87, *p*<0.0001). Follow-up, within-genotype pairwise comparisons indicated that all but the Vip and Sst groups spent significantly more time in the motivated state than in the under-motivated state (*p* < 0.01 for comparisons in Cux2, Rbp4, and Slc17a7 groups).

The consistent progression from over-motivation to under-motivation likely reflects waning engagement due to decreasing thirst in the session. Supporting this, licking reaction times (pooled across mice) were shortest when mice were over-motivated but longest when under-motivated (Figure 4A). Additionally, consumption lick counts (the number of licks in a 5 second window following reward delivery, which is a metric of response vigor) were highest when mice were over-motivated but lowest when under-motivated (Figure 4B) (Berditchevskaia et al., 2016).

**Figure 4.**
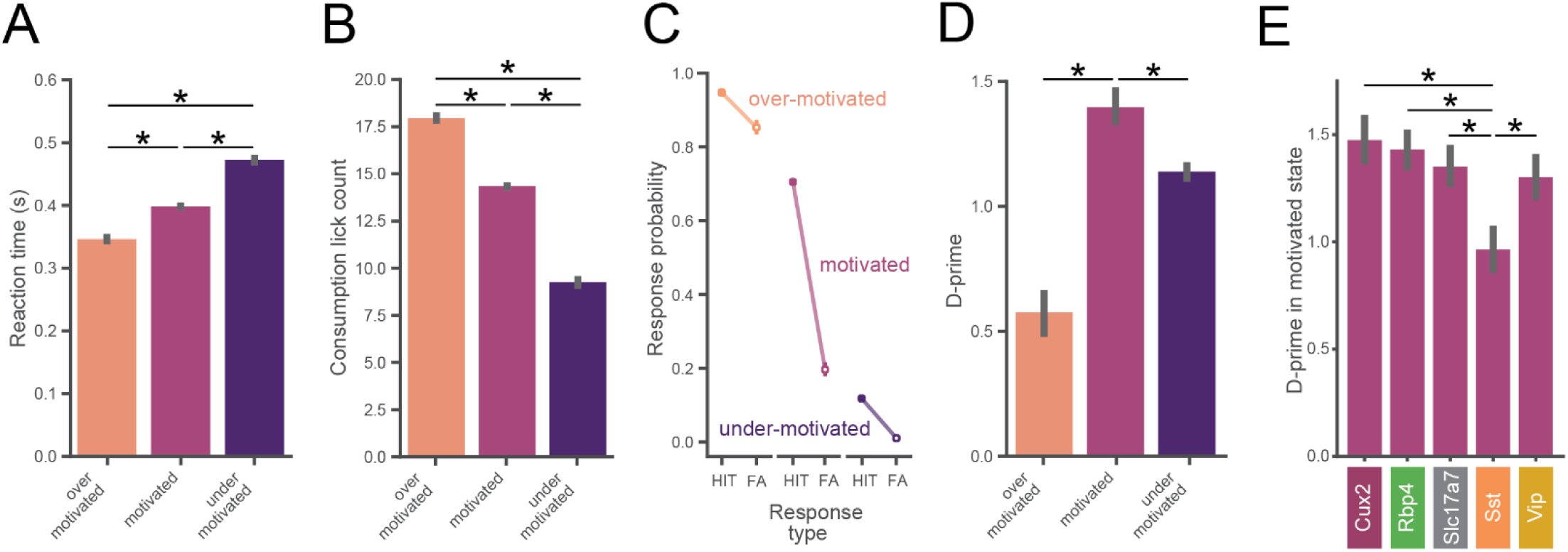
Task performance varies across motivation states. A) Reaction times are slower with lower motivation. In panels A-D, data is pooled across all mice (n=56) and motivation state is defined as in Figure 3E. B) Total number of water consumption licks is less with lower motivation. Total licks are counted in a 5 second window following reward. C) Hit and false alarm rates in each motivation state (defined as in Fig 3E). D) Inverted-U relationship between d-prime and motivational level. D-prime is higher in motivated compared to over- and under-motivated states. E) D-prime in the motivated state for each genotype. The Sst group exhibited a lower d-prime than each of the other genotypes in the motivated state.

### Behavioral performance varies with motivation

The probability of a behavioral response (averaged over all images) varied with motivation levels, as expected from our criterion-based definition (Figure 4C). When over-motivated, both hit and false alarm rates were high. In the more optimal motivational range, hit rates were high but false alarm rates were low. Finally, when under-motivated, mice showed low hit and false alarm rates.

To assess psychophysical performance for each motivational state we computed d-prime values by pooling across all trials from all mice in matched motivational states in order to reduce the impact of epochs with low trial counts (which would provide less accurate estimates of d-prime). We found an inverted-U shape relationship between d-prime and motivation level (Figure 4D), consistent with both classic (Duffy, 1957; Yerkes and Dodson, 1908) and recent studies (Mcginley et al., 2015). We performed a series of pairwise hypothesis tests on the bootstrapped d-prime distributions (Saravanan et al., 2019) and report the bootstrapped probabilities (p_boot_). D-prime was greater in the motivated state than in both the under- and over-motivated states (p_boot_ < 0.001). Thus, periods of ‘optimal’ motivation corresponded to the highest performance as measured with d-prime. Supplemental Figure 3 illustrates how the relationship of d-prime and motivation varies with different criterion thresholds for defining motivational states.

We next computed d-prime values in the motivated state separately for each genotype using the same bootstrap analysis described above. Motivated d-prime values were not significantly different across genotypes, except for the Sst group which had a lower d-prime compared to each of the other groups (p_boot_ < 0.001).

### Highly correlated perception across transgenic lines in motivated state

In the final stage of training (stage 3), mice perform the visual change detection with a set of 8 natural scene images (Figure 5A). In total, mice see 8×8=64 unique image-pair transitions (8 of these are no-change transitions, which define catch trials). On average, mice displayed a range of response probabilities to the 64 unique image pairs, indicating some transitions were more difficult than others (Figure 5B,C). This pattern of responses across image transitions reflects the mice’s perceptual landscape and this might differ across transgenic lines. Thus, we next sought to determine how similar was the pattern of behavioral responses across genotypes and whether this was motivation-dependent.

**Figure 5.**
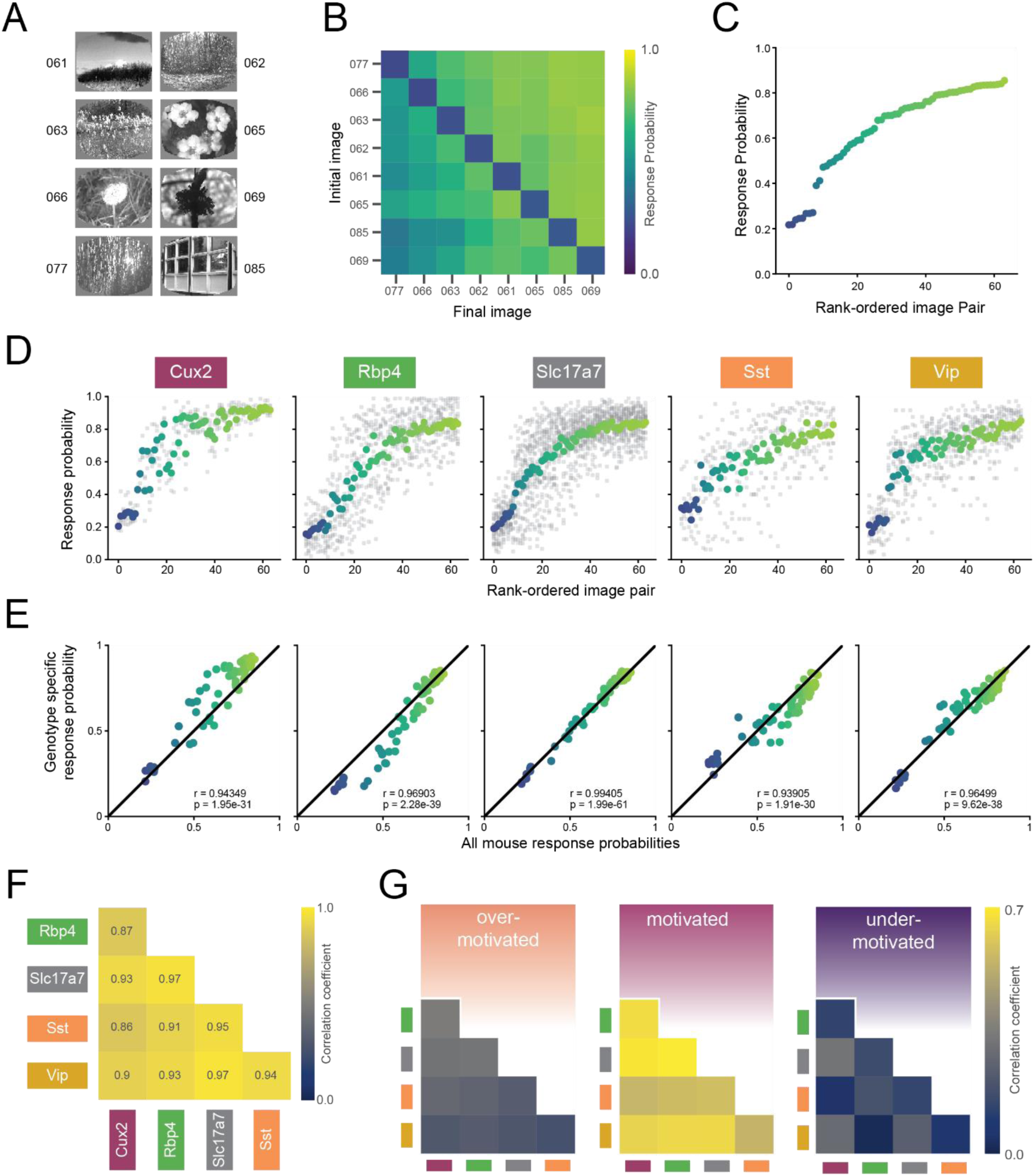
Similar perception across mice during motivated epochs. A) Eight natural scene images used during Stage 3 of change detection task (see Figure 2A). Number indicates label from DeVries et al., 2020. B) Response rate for all pairwise image transitions in the motivated state (average of all mice). C) Average response rate for each image-pair transition. X-axis is ordered by average response rate across all 64 transitions. D) Mean response rate for each image-pair, separated by genotype. The color for each image pair is conserved from (C). Gray points show response rate for each transition for each mouse. E) In the motivated state, each genotype’s pattern of responding was strongly correlated with the average over all mice. F) Response patterns in the motivated state are strongly correlated between all genotypes. G) Bootstrapped correlations of response patterns across genotypes are higher in the motivated compared to under- and over-motivated states. Absolute values are lower compared to (F) due to subsampling to match small trial counts in over-motivated state.

The rank order of the response probabilities for the 64 transitions were largely conserved across genotypes (Figure 5D), and each genotype’s pattern of behavioral responses correlated strongly with the average of all mice (Figure 5E; r-values of 0.93 to 0.99, p-values < 4E-29). Moreover, each transgenic line strongly correlated with the others indicated by significant pairwise correlations between all possible pairs (Figure 5F; *r*-values of 0.82 to 0.97, p-values < 4E-7). To compare the strength of these correlations across the three motivational states, we performed a bootstrapping analysis in which we used subsampling to match sample sizes of each transgenic line across motivational states (see Methods). We found that response correlations were highest in the optimally motivated state compared to over- and under-motivated states for all genotype combinations (Figure 5G, all p-values < 2.1E-164).

## Discussion

We set out to characterize learning and behavioral performance of multiple transgenic mouse lines on a visual change detection task and to further understand how variation in motivation influences performance once trained. Overall, our results show that despite some differences in learning and motivation, the five transgenic mouse lines we tested have highly correlated visual perception during optimally motivated states.

### Standardized behavior training of transgenic mice

An overarching goal of this work is to establish standardized training protocols to implement a robust behavior pipeline for characterization of cellular physiology using our *Allen Brain Observatory*. The transgenic lines we tested allow measurement of activity in specific subsets of excitatory cells (Cux2-CreERT2: Layers 2/3, Rbp4-Cre_KL100: Layer 5, Slc17a7-IRES2-Cre: Layers 1-6), and distinct inhibitory cell classes (Sst-IRES-Cre, Vip-IRES-Cre). As part of our development process it was important to anticipate experimental throughput by quantifying learning times and verifying robust task performance in these transgenic lines. Our results described here extend the basic phenotypic characterization of these transgenic lines (Daigle et al., 2018).

We trained all mice with an automated protocol that applied consistent parameters and task progression rules. All transgenic lines could be reliably trained in several weeks to perform the task using our protocol. Vip mice required significantly longer to reach the final stage of the task but performed at similar levels once trained. Additionally, although Sst mice learned the task quickly, they exhibited lower performance (d-prime) in the motivated state.

Future work can decipher the cause of learning, motivation, and performance differences in these transgenic lines, and whether it relates to neuronal GCaMP6 expression, developmental effects, and/or off-target effects on other brain or body systems. For instance, developmental disruption of Vip interneurons is known to impair perceptual learning in mice (Batista-Brito et al., 2017). Additionally, Sst transgenic mice have an increased incidence of health-related issues including a propensity for dermatitis (Allen Institute for Brain Science, 2016). Differences in task training times have been noted in other transgenic lines such as Vgat-ChR2 mice (Resulaj et al., 2018), which express channelrhodopsin in inhibitory neurons. Importantly, despite differences in learning and motivation, we found that perceptual decisions were very consistent across different lines when comparing matched motivational states.

### Motivation is non-stationary

In most mice, motivation systematically decreased over each behavioral session. This likely represents a decrease in thirst-based motivation as water is consumed in the task. Consistent with this, we observed changes in licking behavior, including lick reaction time (lick latency) and consumption lick count (response vigor), which have been linked to motivational changes (Berditchevskaia et al., 2016). Interestingly, recent work suggests a brain-wide network is involved in thirst regulated motivation (Allen et al., 2019). Thus, characterizing changes in thirst-based motivation will likely be important for interpreting neural activity measurements in tasks involving water reward.

We used a metric from signal detection theory, ‘criterion’ (Green et al., 1966), to estimate motivation and to categorize states in the behavior sessions as over-motivated, motivated, or under-motivated. Future work can develop improved methods for identifying and quantifying behavioral states including generalized linear models and hidden Markov models (Calhoun et al., 2019; Wiltschko et al., 2015). These methods have the potential to provide a more powerful description of motivation, task-engagement, and other latent variables, and might also reduce the need for the temporal binning approach used here. In addition, they could help to explore how task contingencies and reinforcement structures affect motivation state and could provide insight into the factors that shape task learning, behavioral strategy, and ultimate performance levels.

It will be important in future work to relate motivation to other behavioral and physiological states. Pupillometry measurements can reflect internal states including levels of arousal and task-engagement (Mcginley et al., 2015; Vinck et al., 2015). In addition, animal movements, including spontaneous actions and fidgets (Musall et al., 2019; Stringer et al., 2019), can be captured with whole body or face cameras and analysis of these behavioral data streams might provide additional quantitative correlates of motivation.

### Similar perception across transgenic mice

We used our behavioral task to assess natural image change detection in transgenic mice. Expert mice can differentiate each of the unique combinations of natural images tested, although some image pair transitions are more difficult to distinguish than others, consistent with a target/distractor paradigm in mice (Yu et al., 2018). The mouse lines we tested here show correlated behavioral responses, and this correlation is very high when mice are compared under matched motivation states. Thus, these transgenic lines show similar patterns of perception despite some differences in learning rates, motivation dynamics, and d-prime values.

In forthcoming physiological experiments, we will measure neural activity in these mice to characterize cellular correlates of change perception, task-engagement, short-term working memory, and temporal expectation. In an initial study of layer 2/3 excitatory and Vip inhibitory cells in visual cortex, we found that excitatory cells provide selective image coding in the task, whereas Vip cells undergo dramatic changes in activity dynamics with learning (Garrett et al., 2020). Large-scale systematic mapping of activity in different cell classes across the brain will provide insights into how these interactions mediate neural processing to guide behavior and learning.

## Acknowledgements

We thank the Allen Institute founder, Paul G. Allen, for his vision, encouragement and support. We thank Corbett Bennett, Sam Gale, Brian Hu, Jerome Lecoq, Stefan Mihalas, Alex Piet, Nick Ponvert, and Christof Koch for helpful discussions and feedback on the manuscript.

## Author contributions

Conceptualization: S.R.O, P.A.G., D.R.O., S.M., J.K., M.G. Supervision: S.R.O., P.A.G.

Data Collection: L.C., S.C., P.A.G., K.M., J.S.

Investigation, validation, methodology, and formal analyses: D.R.O., S.R.O, P.A.G., S.M., J.K., M.G.

Software: D.W., D.R.O., J.K.

Data Curation: D.R.O., J.K.

Visualization: D.R.O, P.A.G., J.K.

Original draft written by S.R.O, P.A.G., D.R.O with input from J.K.

All co-authors reviewed the manuscript.

## Methods

### Mice

All experiments and procedures were performed in accordance with protocols approved by the Allen Institute Animal Care and Use Committee. Male and female transgenic mice expressing GCaMP6 in various Cre-defined cell populations were used in these experiments (Madisen et al., 2015). Mice were maintained on a reverse 12-hour light cycle (off at 9am, on at 9pm) and all experiments were performed during the dark cycle. [Table of mice used in experiments in Supplemental Table 1].

### Surgery

Headpost and cranial window surgery was performed on healthy mice that ranged in age from 5-12 weeks. Pre-operative injections of dexamethasone (3.2 mg/kg, S.C.) were administered at 12h and 3h before surgery. Mice were initially anesthetized with 5% isoflurane (1-3 min) and placed in a stereotaxic frame (Model# 1900, Kopf, Tujunga, CA), and isoflurane levels were maintained at 1.5-2.5% for surgery. An incision was made to remove skin. The exposed skull was levelled with respect to pitch (bregma-lambda level), roll, and yaw. The stereotax was zeroed at lambda using a custom headframe holder equipped with stylus affixed to a clamp-plate. The stylus was then replaced with the headframe to center the headframe well at 2.8 mm lateral and 1.3 mm anterior to lambda. The headframe was affixed to the skull with white Metabond and once dried, the mouse was placed in a custom clamp to position the skull at a rotated angle of 23° such that visual cortex was horizontal to facilitate the craniotomy. A circular piece of skull 5 mm in diameter was removed, and a durotomy performed. A coverslip stack (two 5 mm and one 7 mm glass coverslip adhered together) was cemented in place with Vetbond (Goldey et al., 2014). Metabond cement was applied around the cranial window inside the well to secure the glass window. Post-surgical brain health was documented using a custom photo-documentation system. One, two, and seven days following surgery mice were assessed for overall health (bright, alert, and responsive), cranial window clarity, and brain health. Upon successful recovery from surgery mice entered into behavioral training.

### Behavior Training

#### Water restriction and habituation

Throughout training mice were water-restricted to motivate learning and performance of behavioral task (Z. V. Guo et al., 2014). Prior to water restriction mice were weighed once daily for three days to obtain a stable, initial baseline weight. During the first week of water restriction mice were habituated to daily handling and increasing durations of head fixation in the behavior enclosure over a five-day period. The first day of behavior training began 10 days of water restriction. Mice were trained 5 days per week (Monday-Friday) and were allowed to earn unlimited water during the daily 1 hour sessions; supplements were provided if earned volume fell below 1.0mL and/or body weight fell under 80-85% of initial baseline weight. On non-training days mice were weighed and received water provision to reach their target weight, but never less than 1.0 mL per day).

#### Apparatus

Mice trained in custom-designed, sound-attenuating behavior enclosures equipped with a 24” gamma-corrected LCD monitor (ASUS, #PA248Q). Mice were head-fixed on a behavior stage with 6.5” running wheel tilted upwards by 10-15 degrees. The center of the visual monitor was placed 15 cm from the eye and visual stimuli were spherically warped to account for the variable distance from the eye toward the periphery of the monitor. Water rewards were delivered using a solenoid (NI Research, #161K011) to deliver a calibrated volume of fluid through a blunted, 17g hypodermic needle (Hamilton) positioned approximately 2-3 mm away from the animal’s mouth.

#### Change detection task

##### Overview

Mice were trained for 1 hour/day, 5 days/week using a behavioral program implementing a go/no-go change detection task schematized in Figure 1. Briefly, mice were trained to lick a reward spout when the identity of a flashed visual stimulus changed identify. If mice responded correctly within a short, post-change response window (115-715ms) a water reward (5-10uL) was delivered. The four stages of the training protocol are shown in Table 1.

| Stage | Stimulus | Stimulus Presentation | Resp. Window (ms) | Contingent Rewards | Duration (min) |
| --- | --- | --- | --- | --- | --- |
| 0 | Square-wave gratings | Static | NA | False | 15 |
| 1 | Square-wave gratings | Static | 1 | True | 60 |
| 2 | Square-wave gratings | 250 ms stimulus; 500 ms gray period | 600 | True | 60 |
| 3 | Natural Images | 250 ms stimulus; 500 ms gray period | 600 | True | 60 |

On Day 1 of the automated training protocol mice received a short, 15-min “open loop” session during which non-contingent water rewards were delivered coincident with 90° changes in orientation of a full-field, static square-wave grating (Stage 0). This session was intended to 1) introduce the mouse to the fluid delivery system and, 2) provide the technician an opportunity to identify the optimal lick spout position for each mouse. Each session thereafter was run in “closed loop”, and progressed through 3 phases of the operant task: 1) static, full-field square wave gratings (oriented at 0° and 90°, with the black/white transition always centered on the screen and the phase chosen randomly on every trial), 2) flashed, full-field square-wave gratings (0° and 90°, with phase as described in 1), and 3) flashed full-field natural scenes (8 natural images used in the *Allen Brain Observatory*).

##### Progression through training stages

Starting with Stage 1 mice were required to achieve a session maximum performance of at least d-prime=2 (calculated over a rolling 100 trial window without trial count correction) during two of the last 3 sessions (Advancement Criteria). The fastest progression from Stage 1 to Stage 3 was 4 training days.

##### Behavior session and trial structure

Each behavior session consisted of a continuous series of trials, schematized in Supplemental Figure 1A. Briefly, prior to the start of each trial a trial-type and change-time were selected. Trial-type was chosen based on predetermined frequencies such that “GO” and “CATCH” trials occurred with predetermined probabilities. In stages 1 and 2, the catch probability was set at 25%, but no more than three consecutive trials of a given type were permitted, leading to an effective catch probability of ∼36%. In stage 3, the catch probability was initially set at 12.5% (given that the 8 same-to-same changes represented 8/64 possible image changes), which, combined with the maximum of 3 consecutive go/catch trial rule, led to an effective catch probability of ∼30%. However, later sessions implemented a matrix sampling algorithm that ensured that each image transition was sampled equally, pushing the actual catch probability to ∼12.5%. Change-times were selected from a truncated exponential distribution ranging from 2.25 to 8.25 seconds (mean of 4.25 seconds) following the start of a trial. Due to computational lag when aligning change-time with a stimulus flash, the actual distribution of change times was shifted to the right by one 750ms flash cycle (with only a small fraction of changes occurring at 2.25 seconds) resulting in a mean change time of 4.2 seconds. In trials when a mouse licked prior to the stimulus change the trial was reset, and a timeout period was imposed. The number of times a trial could be reset before re-drawing the timing parameter was limited to five. In all, this trial structure leads to a sampling of “GO” and “CATCH” trials, that when combined with mouse responding, yields “HIT”, “MISS”, “FALSE ALARM”, and “CORRECT REJECTION” trials.

In addition to the four trial types described above, behavior sessions contained a subset of “free reward” trials (“GO” trials followed immediately by delivery of a non-contingent reward). Behavior sessions across all phases began with 5 “free-reward” trials. Additionally, in order to promote continued task performance throughout the behavior session in a subset of sessions “free reward” trials were delivered after 10 consecutive “MISS” trials.

### Data analysis

Analysis was performed using custom scripts written in Python v3.7.5 (including Pandas v0.24.2, Numpy v1.16.4, Scipy v1.3.2 and Statsmodels v0.10.1) and GraphPad Prism (v8.0.1). Plots were generated using Matplotlib v3.1.1 and Seaborn v0.9.0.

Behavioral performance was quantified with the signal detection metrics of d-prime and criterion, which are both a function of hit and false alarm rates.

#### Hit and false alarm rates

The hit rate was calculated as the fraction of go-trials in which the mouse licked in a 0.115 to 0.715 second window following the display-lag-compensated image display time. Catch trials were defined as trials in which there was no image change. However, for calculation of the false alarm rate, a response window was defined following one of the flashes using the same statistics as in the go trials. False alarm rates were calculated as the fraction of catch-trials in which animal emitted a lick in this response window. Unless otherwise noted, hit and false alarm rates were corrected to account for trial counts using the following formula (Macmillan and Creelman, 2004):

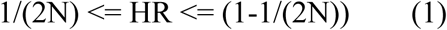

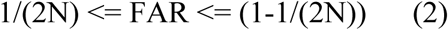

Where HR and FAR represent the hit and false alarm rates, and N represents the number of the respective trial type.

#### D-prime (d′)

d-prime, which is a measure of the relative difference in response probabilities across the two trial types, is defined as:

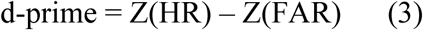

in which Z represents the inverse cumulative normal distribution function.

#### Criterion

Criterion, which is a measure of the underlying bias of the subject to emit a response, is defined as:

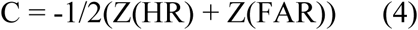

Criterion therefore varies from negative values for high response biases (high hit and false alarm rates) to positive numbers for low response biases (low hit and false alarm rates). In general, our figures represent criterion with the sign inverted, thus mapping states of low motivation to negative criterion values and states of high motivation to positive criterion values.

In Figure 1C, trials were pooled across all included sessions and all licks occurring within 1.5 seconds of the stimulus display time (approximately two full stimulus flash cycles) were included. The cumulative distributions were calculated after grouping trials by animal ID, with each green line representing one animal’s cumulative distribution of licks on go trials. The dark black line represents lick times pooled over all trials and mice.

The max d-prime plotted in Figure 2A represents the peak values calculated from a 100-trial rolling window without trial count correction and represent the actual values used when calculating advancement criteria in the automated training algorithm. Median reaction time (RT) is calculated over the entire duration of the session.

Figure 2B represents the training stage for each of the 60 mice in the dataset. White values are missing data due to animals being removed from the study (for health- and non-health-related causes) prior to the 15 days displayed in the plot.

Figure 2C represents the number of training days to reach stage 2 (light hues) and stage 3 (dark hues) for each genotype. The error bars represent the 95% bootstrapped confidence interval for all mice that reached stage 3 in each genotype. A main effect of genotype on training time was identified using the Kruskal-Wallis H-test for independent samples. Pairwise post-hoc Dunn’s multiple comparisons tests were used to identify significant training time differences between groups.

The engagement plots in Figure 3B represent criterion as described in eq. 4, calculated without trial count correction and with the sign inverted to represent higher states of motivation in the positive direction. The light gray lines represent criterion values calculated in 10 minute time bins for each of the sessions performed by that mouse. The black line represents the mean value across all sessions in each 10 minute bin, with error bars representing standard deviation.

In Figure 3C, each light gray line represents the criterion values traversed by a single mouse in a single session (same as in 3B), with every session from every mouse shown. The black line represents the average across all sessions. Error bars represent standard deviation.

In Figure 3D, every row in the matrix represents one mouse, with each cell representing the criterion value for that mouse in a given 10 minute epoch, averaged across all expert-level sessions that the mouse performed. Colors range from dark (low criterion, low motivation) to light (high criterion, high motivation). Mice are grouped by genotype, and by average criterion value within genotype.

The histogram in Figure 3E shows the range of criterion values assigned to every 10 minute epoch across all 1100 analyzed sessions, regardless of mouse or genotype. Epochs without at least one hit trial and one false alarm trial (1.2% of the total) were not assigned a criterion value (and thus not included). The sign of the criterion metric is inverted so that low motivation states (high criterion values) lie to the left. Thresholds were drawn at criterion values of 1.25 and −1.25 with every 10 minute epoch being assigned a label of ‘motivated’ (73.2%), ‘under motivated’ (18.6%) or ‘over motivated’ (6.9%) depending on the criterion value in that epoch.

In Figure 3F, each bar represents the average fraction of time spent in a given motivation state for all animals of a given genotype. Error bars represent bootstrapped 95% confidence intervals.

All data shown in Figure 4 relies on individual trial data separated by the assigned motivation state, as described in Figure 3E. Trials in expert-level sessions were given a label (either ‘motivated’, ‘under motivated’ or ‘over motivated’) based on the label of the 10 minute epoch in which they occurred. Trials that occurred in epochs without a label (i.e., epochs without at least one hit trial and one false alarm trial) were excluded from the analysis.

Figures 4A and 4B include pooled data from mice in all expert sessions. In Figure 4A, reaction time is calculated as the time to first lick in all hit trials in each of the three motivation states. In Figure 4B, reward lick count is calculated as the total number of licks in a 5 second window following reward delivery on every hit trial. Pairwise independent t-tests were used to assess significance metrics across motivation states.

Figures 4C and 4D used performance data calculated for each mouse in each of the three motivation states, with Figure 4E using data only from the motivated state. In Fig. 4C, hit and false alarm rates were calculated by pooling across all trials in a given motivation state. Data points represent mean response probabilities (hit or false alarm rates) from the pooled data. Error bars representing 95% confidence intervals after a 1000 iteration bootstrap procedure in which N trials from each motivation state were sampled with replacement, with N set to the lowest trial count in any of the three motivation states (9382 trials).

The d-prime values displayed in Figures 4D and 4E are derived directly from the hit and false alarm rates shown in Figure 4C, with the value recalculated on the output of every bootstrap iteration. Error bars representing 95% confidence intervals on the bootstrapped d-prime values in each state. Statistical comparisons were performed by calculating the total density of the joint probability distribution on one side of the unity line, yielding a probability, p_boot_, that null hypothesis is true (Saravanan et al., 2019). Pairwise comparisons were deemed significant if the fraction of overlap was less than the Bonferroni corrected two-tailed alpha. The resolution of p_boot_ was limited by the number of bootstrap iterations (1000), providing a minimum measurable value of 0.001.

The grand-average response matrix in Figure 5B represents the probability of response for each image pair, with catch trials (same-to-same transitions) on the diagonal. The matrix was calculated by first calculating a response matrix for each mouse (using all trials in the motivated state), then averaging together matrices across all mice in a given genotype, and finally averaging together matrices across all five genotypes. Thus, mice with different trial numbers will contribute equally to the genotype averages, and genotypes with different mouse numbers will contribute equally to the grand-average. Of the 56 mice with at least one expert session, four mice with fewer than an average of 4 presentations of each of the 64 possible natural image pairs (256 total trials) were excluded from these and subsequent analyses.

Figure 5C represents the data in Figure 5B, but with values from the matrix unwrapped into vector form, then rank-sorted by response probability. The color value of each dot matches the color value of the corresponding square in Figure 5B. Gray dots represent each of the five genotype averaged response probabilities for the corresponding image transition pair.

The response probability curves in Figure 5D represent the response probabilities for each of the 64 image combinations for each of the five genotypes, with the rank order from Figure 5C preserved. Each gray dot represents the response probability for an individual animal for a given image pair. The plots in Figure 5E represent the correlation between the genotype-averaged response vector and the grand-average response vector in Figure 5C, with *r* and *p*-values representing Pearson correlation coefficients.

Figure 5F shows the Pearson correlation coefficients for each pairwise combination of genotype response vectors (from Figure 5D). Diagonal terms (with p = 1.0) and above diagonal terms (with values equal to the below diagonal terms) are excluded from the display.

To compare correlation values across the motivational states, we performed a bootstrap analysis in which we first pooled all trials from a given genotype, then subsampled trials with replacement from each genotype using the smallest trial count from any genotype/motivation state combination (708 trials for the Rbp4-Cre mice in the over motivated state). New response matrices were calculated on the subsampled data and Pearson’s correlation coefficients were calculated for each pair of genotypes. The process was then repeated 1000 times. Correlation coefficients shown in Figure 5G represent the mean values across all iterations. The range of correlation values from the bootstrap process were compared in the over motivated vs. motivated and the under motivated vs. motivated conditions for each pairwise combination of genotypes using Wilcoxon signed-rank tests.

---

**Supplemental Figure 1.**
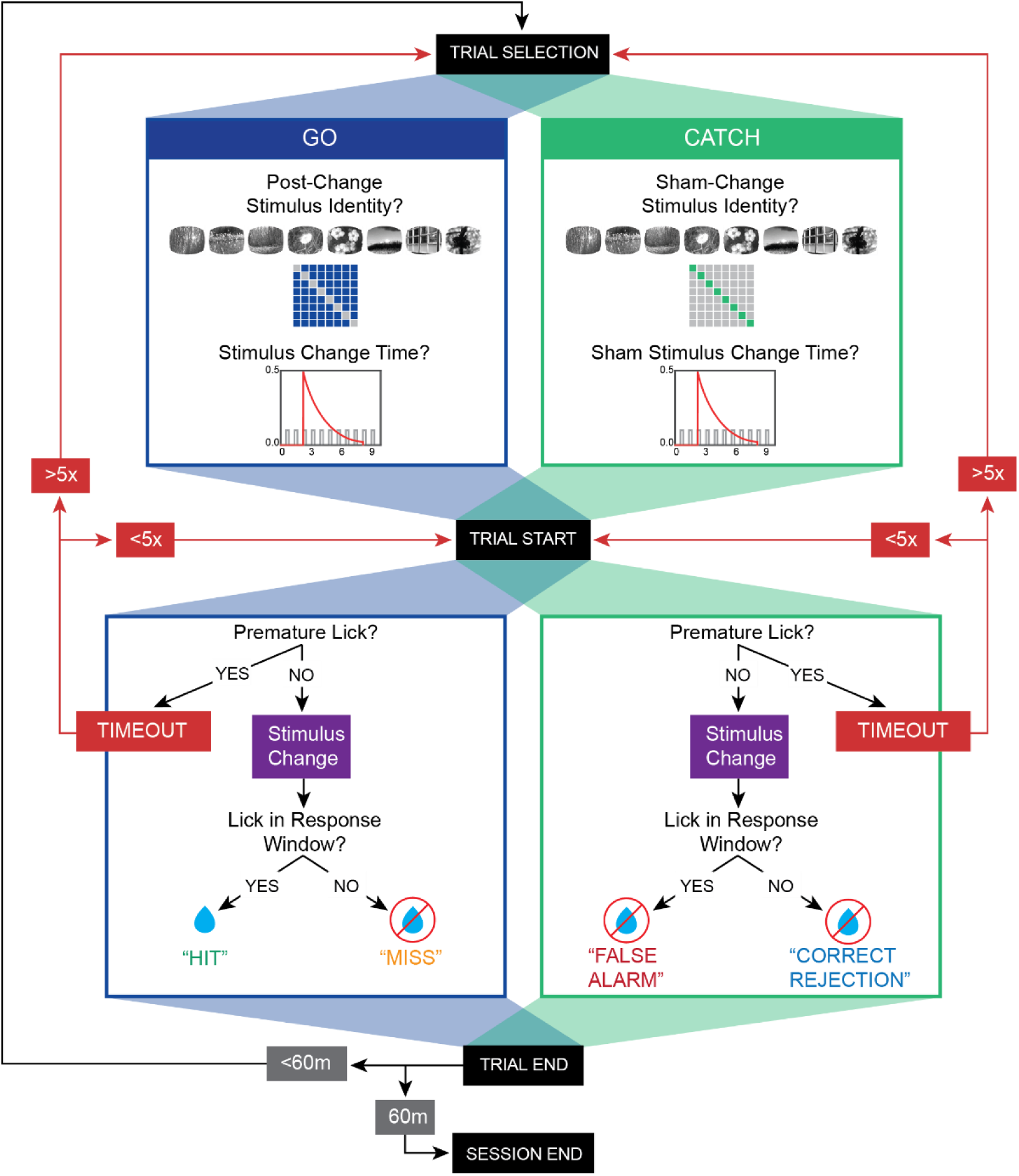
Task flow diagram. The “Flashed Images” stage of the change detection task consists of 8 images, resulting in 64 possible image transitions, including both GO and CATCH trials. GO trials comprise 87.5% of all trials and are represented in the off-diagonal portions of the 8×8 change matrix. CATCH trials comprise 12.5% of all trials and are represented in the diagonal of the matrix. Each trial was first selected as either GO or CATCH and a post-change (or sham-change) image identity was chosen from the change matrix. The stimulus change (or sham-change) time was then selected from a truncated exponential distribution between 2.25s to 8.25s. As stimuli are presented every 715 ms, the actual change time was determined as the nearest flash from the drawn time. Once a trial started, a premature lick (i.e., a lick that occurred prior to the predetermined change time) resulted in a timeout and the trial was restarted. If an animal caused a trial to timeout 5 times, a new trial was selected. If no premature licks were recorded, the trial progressed and the stimulus change occurred at the predetermined change-time. On GO trials, a lick detected within 600ms response window resulted in a “HIT” (and subsequent reward delivered) whereas a lack of response resulted in a “MISS”. On CATCH trials, a lick within the window following the sham-change resulted in a “FALSE ALARM”, whereas a lack of response resulted in a “CORRECT REJECTION”. Following the stimulus change and response window the trial ended and a new trial was selected. The session ended after 60 minutes.

**Supplemental Figure 2.**
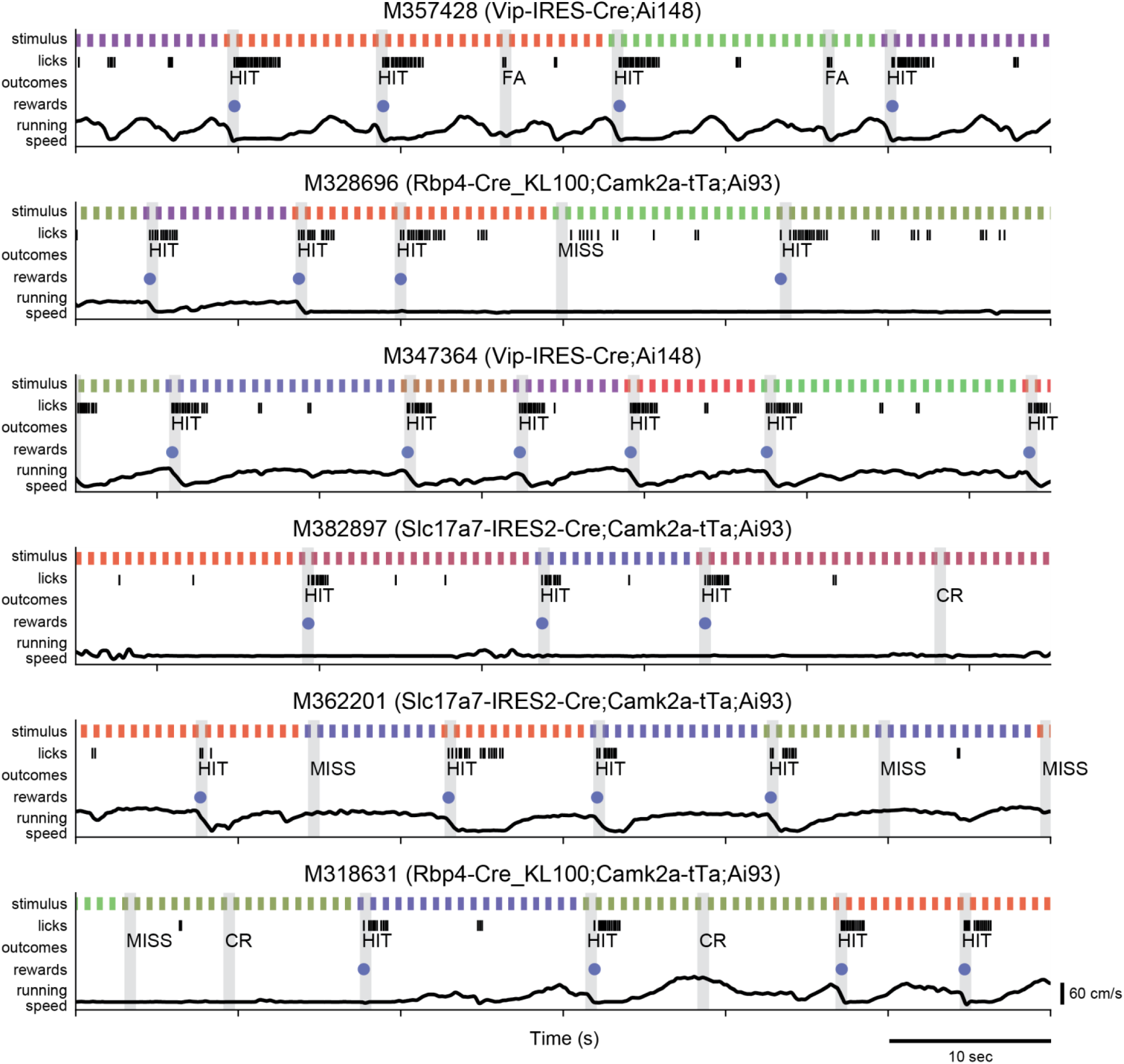
Example behavioral segments. One-minute examples of behavior from 6 mice of various genotypes. Each example shows stimulus (color-coded by image identity to show when changes occur), licks, trial outcome, reward delivery, and running speed.

**Supplemental Figure 3.**
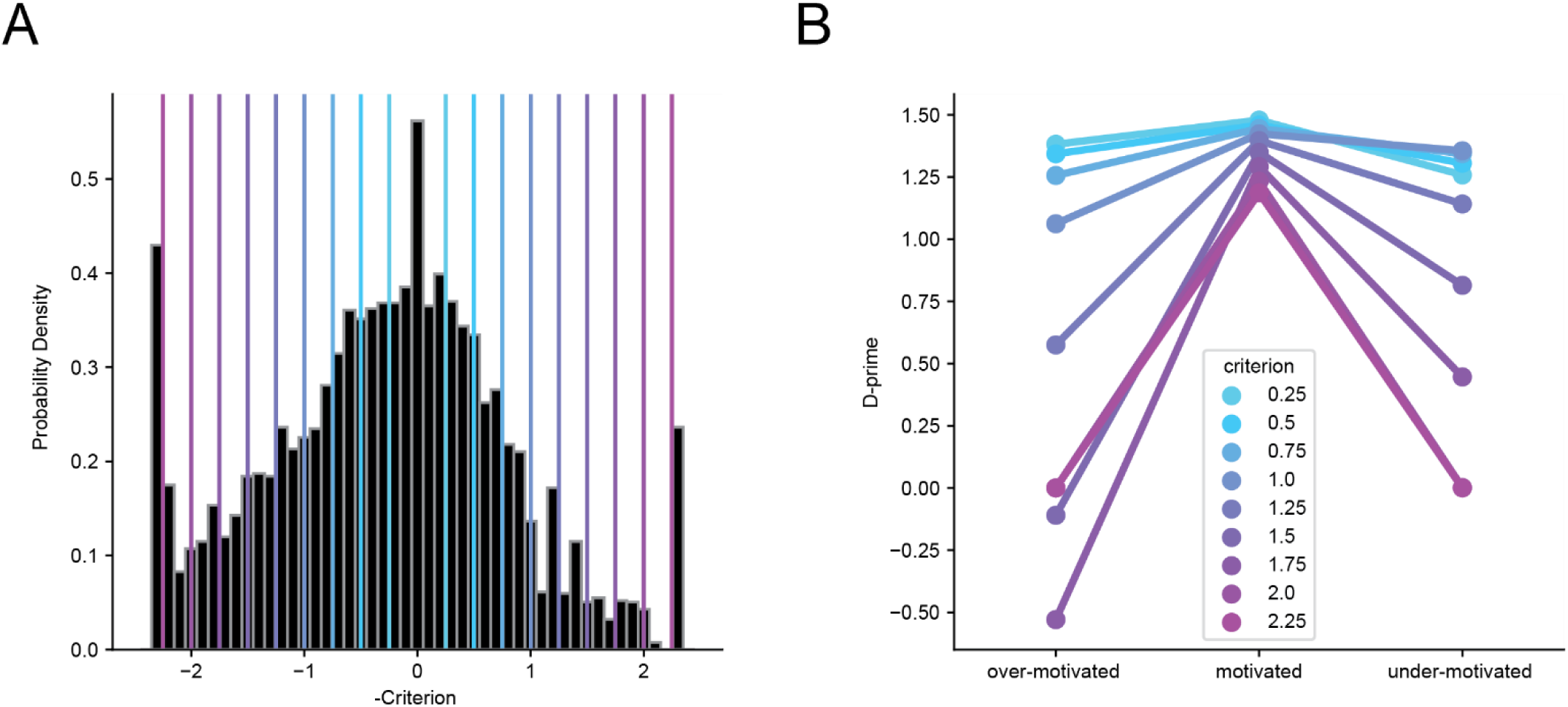
D-prime in each motivation state for a range of criterion thresholds. A) The histogram of criterion values (same as Fig 3E) with a range of criterion thresholds drawn. Thresholds range from +/- 0.5 to +/- 2.25 in increments of 0.25. In every case, the ‘motivated’ epochs are designated as those that fall between the thresholds, the over motivated epochs are those that fall to the right of the higher threshold and the under motivated epochs are those that fall to the left of the lower threshold. Note that thresholds of +/- 1.25 were used in the main figures. B) D-prime calculated on all pooled trials in each of the three motivation states for the range of criterion values shown in A.

**Supplemental Table 1.**
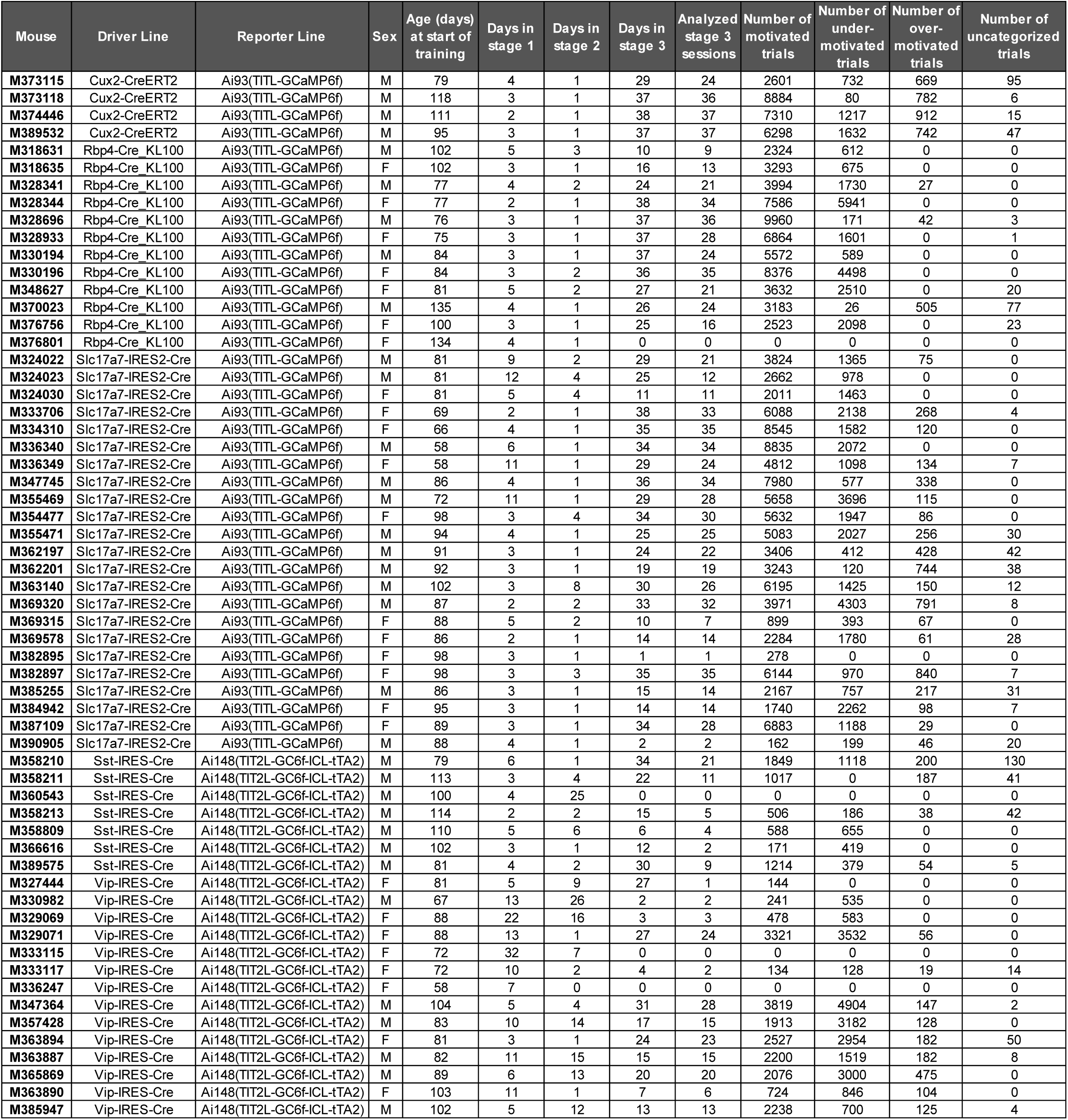
Mice included in study.

